# Beyond single markers: bacterial synergies identified by Multidimensional Feature Selection reveal conserved microbiome disease signatures

**DOI:** 10.64898/2026.04.13.718216

**Authors:** Kinga Zielińska, Witold Rudnicki, Paweł P Łabaj

## Abstract

The gut microbiome encodes disease-relevant information not only in the abundance of individual taxa and functions, but in the way they co-occur and interact. Yet metagenomic analyses have largely relied on univariate approaches that evaluate features in isolation, systematically overlooking the combinatorial signals that arise from meaningful, disease-informative patterns of microbial co-occurrence. Here, we introduce a framework based on the Multidimensional Feature Selection (MDFS) algorithm to identify synergistic feature pairs - combinations of taxa and functions in which the information carried by a feature significantly increases in the presence of a partner. This includes features which carry no detectable individual information and may therefore be overlooked by univariate analyses.

We first validated the approach on a meta-analysis of colorectal cancer (CRC) cohorts - one of the most competitive microbiome classification benchmarks available - using a leave-one-cohort-out cross-validation framework. Classifiers built solely on our synergy-derived features achieved performance comparable to the state-of-the-art benchmark (AUC = 0.84 vs 0.85). This confirmed that we were able to capture an informative disease signal while revealing microbial interactions not provided by univariate methods. A subset of the pairs was repeatedly selected by the models during feature selection, which indicated their stability. Post-selection effect size analysis revealed their consistent differences between cases and controls.

To learn more about diseases beyond CRC, we extended our framework to 13 additional datasets spanning inflammatory bowel disease, cardiometabolic disease, liver cirrhosis and atherosclerotic cardiovascular disease. We identified 80,790 synergistic pairs in eight of them as well as a set of candidate markers recurring across independent cohorts. Overall, synergistic pairs carried substantially more information about disease status than their individual constituents. Our framework therefore established meaningful microbial co-occurrence patterns which would not be possible with univariate methods alone. The framework is freely available at https://github.com/Kizielins/MDFS_synergies.

**Importance:** Most microbiome studies search for individual gut bacterial species associated with disease. However, bacteria do not act in isolation, and their combined presence or relative balance may be far more informative than any single microbe considered alone. This study presents a computational framework which identifies pairs of gut microorganisms whose co-occurrence or relative abundance carries significantly more information about disease status than their importance alone. Applied to stool metagenomic data from patients with colorectal cancer, as well as individuals with other conditions, we demonstrate that these synergistic interactions are: 1) widespread; 2) reproducible across independent patient cohorts; 3) and resolve disease-relevant microbial relationships which single-feature analyses do not report. Our framework offers a more complete view of how the gut microbiome is altered in disease and provides a principled basis for identifying robust, interaction-based biomarkers.

## Introduction

The human gut microbiome has emerged as a powerful window into host health, with growing evidence that its composition encodes diagnostically and prognostically relevant information across a wide range of diseases (Gilbert et al., 2018; Lynch and Pedersen, 2016). A major thrust of current research has therefore been to identify specific bacterial taxa or functional pathways associated with a given phenotype, most often a health or disease state. The dominant analytical paradigm, whether differential abundance testing, univariate feature selection or single-taxon-versus-outcome modelling, considers features in isolation. Pooled analyses of thousands of metagenomes have, for example, yielded taxonomic health indices such as the Gut Microbiome Wellness Index 2 (Chang et al., 2024), which scores a sample by the presence of individual health- or disease-associated taxa. Our own function- and interaction-based health index Q2PD (Zielińska et al., 2025) instead takes feature relationships into account.

Considering features in isolation is biologically questionable, because members of an ecosystem rarely act independently. Bacteria form functional guilds - groups exhibiting consistent co-abundant behaviour based on availability of resources - and engage in more complex interactions (Wu et al., 2021). Higher-order dependencies, such as competition, generate emergent community-level phenotypes invisible to any single-feature analysis (Ludington and William, 2022). Moving from association to causality also remains an elusive step for designing microbiome-based interventions (Stein et al., 2013; Vos et al., 2022).

Several multivariate methods do consider relationships between features: sparse linear and partial-least-squares models (Lê Cao et al., 2011), compositional balance and log-contrast approaches (Rivera-Pinto et al., 2018; Gordon-Rodriguez et al., 2021; Calle et al., 2023), and tree-based or other machine-learning classifiers, which all combine features and can capture non-additive effects. However, they are optimized for prediction and do not report which features are jointly informative, in which form, and with which partner. Co-abundance network strategies do report interacting species, but they are built on taxa already identified at the individual level. For example, in Derosa et al. (2024) the clusters of interacting species were assembled from individually identified taxa. Such an approach cannot detect pairs whose members are weak on their own and would be discarded as noise. We were therefore missing a way to identify, without pre-selecting on individual signal, which feature pairs were jointly informative and in which form. The relevance of taking feature relationships into account, shown in our work on Q2PD, as well as dual feature importance, motivated us to develop this approach further in the present study.

Here, we present a novel analytical framework which uses the Multidimensional Feature Selection algorithm (MDFS; Mnich and Rudnicki, 2020; Piliszek et al., 2019) in its two-dimensional mode (MDFS-2D). This allows the identification of synergistic feature pairs in metagenomic data. MDFS-2D computes information gain (IG) about the outcome for every pair of features considered jointly, which makes the feature pair the basic unit of our microbiome-disease analysis. Because pairs are evaluated jointly from the start, they are found even when a member would be discarded as noise in a univariate screen. We call a pair synergistic when the IG of the two features together is significantly larger than the IG of the base feature alone (formal definition in Results and Methods). For simplicity and clarity, we restricted our analysis to pairs, although MDFS can evaluate higher-order tuples as well (the implementation supports up to five dimensions). Ultimately, we thus add a new dimension of microbiome structure previously undetected or rejected as noise.

To validate our approach, we tested whether the identified pairs carried predictive value for metagenome-based disease classification. Classification was not our primary objective but served as a validation step. We deliberately chose one of the most competitive benchmarks available, and one of the most extensively meta-analyzed microbiome-disease associations to date (Piccinno et al., 2025): colorectal cancer (CRC), a disease of exceptional clinical relevance (Siegel et al., 2026). The benchmark dataset came with a specialized model reporting one of the highest AUCs to date (AUC = 0.85).

In the validation, we showed that classifiers built on our synergistic pairs achieved performance comparable to the state-of-the-art model, additionally resolving biologically interpretable feature synergies. This included both taxa and functions whose individual signal was weak - and the form in which they were jointly informative (joint enrichment or relative balance). Extending the framework to 13 additional disease cohorts, we identified from a single pair to tens of thousands of synergistic pairs per cohort, with a set of pairs replicating independently across two inflammatory bowel disease cohorts. Together, our results establish pairwise feature synergy as a valid signal in metagenomic data, and show that a part of the microbiome’s disease-relevant structure is resolved only when features are considered in combination.

## Results

### Synergy-oriented framework identifies disease-informative feature pairs

To capture synergistic interactions between microbiome features, we employed the Multidimensional Feature Selection (MDFS) algorithm (Mnich and Rudnicki, 2020; Piliszek et al., 2019). A feature that is uninformative on its own can become informative in combination with a partner, and a univariate analysis cannot identify such a case. MDFS quantifies how informative features are about the label prediction using information gain (IG). This model-free measure derived from mutual information expresses how much knowing a feature’s value reduces uncertainty about the outcome. In 1D mode, MDFS scores each feature on its own, which is equivalent to a standard univariate approach. In 2D, it exhaustively scores every pair of features jointly. Figure 1a presents an overview of the idea.

**Figure 1.**
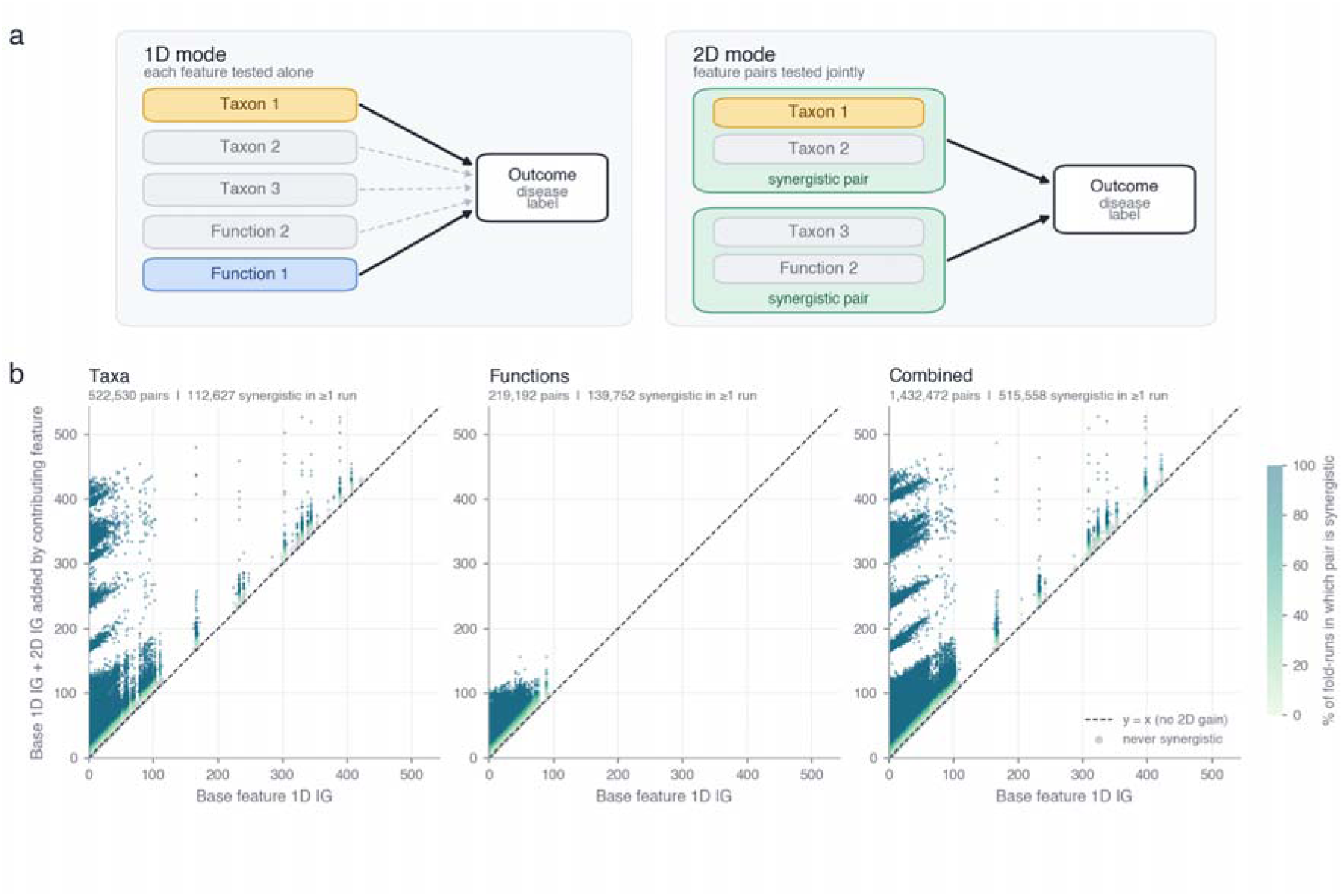
a) Schematic representation of the MDFS framework in 1D and 2D modes. In 1D, each feature is evaluated independently against the outcome, identifying only Taxon 1 and Function 1 as significant. In 2D, all feature pairs are tested jointly. This allows for the identification of features which are not informative by themselves yet are identified as relevant in combination with partners, in the form of synergistic pairs - Taxon 1 + Taxon 2 and Taxon 3 + Function 2. b) Joint 2D information gain of every directed feature pair (base + contributing feature) in the taxonomic (522,530 pairs), functional (219,192) and combined (1,432,472) feature spaces. The x-axis represented the 1D IG of the base feature, while the y-axis is the 1D IG of the base feature plus the IG added by the contributing feature, both averaged over the leave-one-cohort-out folds and MDFS runs (14 training folds × 10 cross validations) in which both features were present. Points on the diagonal carry no 2D gain; the strongest pairs retained for synthetic-feature construction (top 2% by total IG in >= 6 of 10 runs) are those with the largest vertical distance from it.

For each pair, MDFS reports both directions: one feature acts as the base feature and the other as the contributing feature. The joint IG of the pair equals the 1D IG of the base feature plus the IG added by the contributing feature. We call a pair synergistic when this added IG is statistically significant (chi-squared test, Bonferroni-corrected over all feature pairs; see Methods). In other words, the base feature becomes significantly more informative about the disease in the presence of its partner. The definition is directional, so it describes the gain of the base feature given the contributing feature, and the same pair can qualify in both directions.

Synergy in this sense is a statement about a jointly informative co-occurrence pattern. It does not imply a causal or ecological relationship between the two features. We hypothesized that this 2D approach would retain competitive classification performance while revealing biologically interpretable feature relationships that conventional single-feature analyses cannot access.

To validate our hypothesis, we applied MDFS to a well-characterized meta-analysis dataset comprising 17 cohorts of healthy individuals and colorectal cancer (CRC) patients - one of the most extensively benchmarked disease contexts in microbiome research, chosen deliberately for its competitive classification landscape. We implemented a Leave-One-Cohort-Out (LOCO) cross-validation framework: each test-eligible cohort (14 of 17; see Methods) was held out in turn as an independent test set, and MDFS was applied to the remaining cohorts. Pairs were retained by consensus over ten MDFS runs with independent random seeds (see Methods), which gave reproducible sets across two disjoint blocks of ten seeds (Jaccard index 0.95 for taxa and 0.98 for the combined space; Supplementary Table 1). All CRC analyses presented below - MDFS pair detection, synthetic feature construction and random forest classification - were performed within the LOCO framework. To ensure no data leakage, MDFS and all subsequent feature selection steps were applied exclusively to the training cohorts in each fold, and the held-out cohort was used only for evaluation.

Across all three feature spaces examined - taxonomic, functional and combined (see Methods for feature construction) - the total IG of the retained pairs exceeded the 1D IG of their base features, as the significance and synergy filters require. Figure 1b shows the size of this gain for every evaluated pair, relative to its base feature.

To identify the most informative synergistic pairs, we focused on feature combinations where the gain in 2D IG over 1D IG was greatest - that is, pairs located furthest above the diagonal in Figure 1b. Among 10 unordered pairs with the highest mean total IG in each feature space (the 30 highest-ranking directed pairs over all feature spaces are listed in Supplementary Table 2), Fusobacterium nucleatum appeared as either the base or the contributing feature in 9 of them (in taxonomic and combined data). This taxon has been previously associated with CRC, including the benchmark dataset referred to in this study (Piccinno et al., 2025). The strongest partners of F.nucleatum appeared to be P.stomatis, P.micra and D.pneumosintes - all anaerobes with established presence in the oral cavity and subgingival plaque. A fourth high-IG partner, A.obesiensis, is less strictly oral but is similarly associated with polymicrobial anaerobic communities. Pairs of oral anaerobes were consistently the most dominant across folds and MDFS runs. Out of 140 fold-run combinations (14 folds x 10 MDFS runs), in taxonomy-based analysis 4 pairs of oral-associated species were selected in all combinations (including F.nucleatum, P.micra, P.stomatis and D.pneumosintes) and 32 in combined analysis (with G.adiacens, P.stomatis, A.obesiensis, G.morbillorum, D.pneumosintes + and C4 photosynthetic carbon assimilation cycle, PEPCK type (PWY-7117) pathway). These findings are consistent with the benchmark study, which identified oral bacteria as a defining CRC microbiome signature (Piccinno et al., 2025), and serve as a positive control for the framework. A full list of such pairs can be found in Supplementary Table 2.

At the level of individual features, the 2D search subsumed univariate selection, as expected. This is because a feature that is relevant in 1D is also relevant in 2D. In the training set of each fold, MDFS-2D recovered 99-100% of the features selected by MDFS-1D, 93-98% of those selected by Welch’s t-test, and 96-99% of those selected by the Mann-Whitney U test. Recovery was lower for LASSO (69-83%) and ReliefF (66-88%)(Supplementary Figure 1). The reverse did not hold. MDFS-2D selected 1.6-1.9 times more features than these methods, and each of them covered only 53-84% of the MDFS-2D set.

To separate the effect of set size from the effect of pairwise information, we compared MDFS-2D with a size-matched univariate reference: the N features with the highest 1D information gain (IG), where N is the number of MDFS-2D features in that fold. This reference contained 81% of the MDFS-2D features for taxa, 94% for functions and 88% for the combined set. The remaining features (46, 24 and 75 per fold, respectively) were selected because of their pairwise rather than their individual signal. Thus, the MDFS-2D selection is not simply a larger univariate set. A subset of its features (19% of taxa, 6% of functions and 12% of the combined) would not have been selected by the 1D mode from a dataset at the same set size.

Collectively, the prevalence of oral anaerobes among the top synergistic pairs indicates that, in these data, meaningful co-occurrence patterns among oral species carry more information about CRC status than any of these taxa considered individually.

### Synergy-derived features retain benchmark-level classification performance

Identified base features with high 2D IG are particularly relevant if they carry genuine predictive value. We therefore asked whether the MDFS-selected pairs could, when encoded as synthetic features, contribute to competitive disease classification. To confirm this, we integrated them as additional features (further referred to as “synthetic features”: for each MDFS-selected pair, we computed the log-ratio (LR) and the geometric mean (GM) of the two constituent features; see Methods) into a random forest (RF) classification framework using a leave-one-cohort-out (LOCO) cross-validation approach. We compared three modes: univariate features (raw, without any feature selection), synthetic features (constructed based on MDFS 2D pairs with significant IG), and a combined model with both sets. Taxonomic data yielded the highest predictive performance, with all three feature configurations achieving a mean AUC of 0.84-0.85 (raw 0.847, synthetic-only 0.839, combined 0.837) and therefore closely following the state-of-the-art benchmark for these cohorts (0.85). Models utilizing integrated taxonomic and functional data were nearly as accurate, with all three configurations performing on par (AUC = 0.82-0.83; raw 0.825, synthetic-only 0.824, combined 0.827). Conversely, functional features alone provided the lowest discriminative power (AUC = 0.67; raw 0.674, synthetic-only 0.667, combined 0.672). Notably, model performance exhibited cohort-specific variability, with the optimal input type (taxonomy vs. function) and feature mode (univariate vs. synergistic) differing across individual studies (Supplementary Figures 2&3). We note that these results did not increase discriminative performance in this benchmark. Their value, however, was instead in showing that a compact, pair-based representation could retain the disease-relevant information of the full feature space while making the underlying feature dependencies explicit.

In a LOCO validation, the held-out cohort was never seen during training but the remaining cohorts still shaped feature selection and the classifier. Three cohorts were never held out and entered every training set. This was because they did not fulfill pre-defined criteria (two had only CRC samples and one had too few samples, see Methods). In such cohorts, cohort identity is almost perfectly confounded with disease label, which could bias feature selection and the stable-pair lists. To test this, we repeated the entire LOCO pipeline with these three cohorts removed from all training sets (sensitivity run). Classification performance in the taxonomic and combined spaces was essentially unchanged. Mean taxonomic AUC for the raw / synthetic-only / combined configurations was 0.845 / 0.836 / 0.831 in the sensitivity run versus 0.847 / 0.839 / 0.837 in the main analysis; in the combined space it was 0.827 / 0.826 / 0.827 versus 0.825 / 0.824 / 0.827. Functional AUCs increased by about 0.04 (0.714 / 0.709 /0.711 versus 0.674 / 0.667 / 0.672; Supplementary Table 3). The hub structure described below was preserved, with fewer stable combined-model pairs for *P. micra* and *F.nucleatum* (10 and 84 versus 18 and 125 in the main analysis). The identity of individual stable pairs, however, depended on training-set composition: 61 of the 169 stable combined-model pairs of the main analysis were stable in the sensitivity run.

Having confirmed that synergy-derived features support benchmark-level classification performance, we next asked which specific pairs the RF model relied upon most consistently - and whether their frequency reflected genuine biological signal rather than model-specific artefact. A synthetic pair was considered stable if it was selected in at least 80% of LOCO folds and was retained only if its frequency exceeded both constituent features individually. Of the 235 pairs meeting these criteria (226 distinct pairs; nine were stable in two feature spaces), the combined model contributed 169 (taxonomic 40, functional 26) - confirming that cross-domain pairing (taxonomy with function) is particularly effective at capturing latent interactions. In addition, for 207 of the 235 pairs at least one constituent feature had 0% individual frequency, meaning the feature was never selected by the classifier as a raw feature but was consistently retained as part of the pair. Because raw and synthetic features competed for selection in the combined model, this indicated that the pair-level representation was preferred over the individual feature, not that the individual feature lacked disease signal.

Hub analysis revealed a striking pattern: *P. micra*, which was never selected individually in the combined model (0% frequency), appeared in 18 stable pairs of the combined model, as the base (lower-frequency) feature in 13 of them - indicating that, in the combined model, its signal was captured preferentially through its synthetic pair features. This should not be interpreted as an absence of individual signal: *P.micra* is a well-established individual CRC marker (Piccinno et al., 2025), and its 0% raw selection frequency most likely reflected competition with the synthetic features derived from it. Indeed, *P. micra* was individually significant in 1D MDFS in 14 of the 14 LOCO folds for the combined feature set and in 14 of 14 for the taxonomic feature set (every MDFS run of those folds; Methods). Synthetic features are by construction correlated with their constituents, so the preference of the classifier for a pair-level feature over a raw one reflected a choice between largely redundant representations, not an absence of individual signal. Conversely, *F.nucleatum* (64% individual frequency) acted predominantly as a contributor, providing IG to 123 of its 125 stable partner features (Figure 2a). This pattern shows that, when meaningful co-occurrence is explicitly encoded, the classifier consistently prefers pair-level representations of these taxa over their individual abundances.

**Figure 2.**
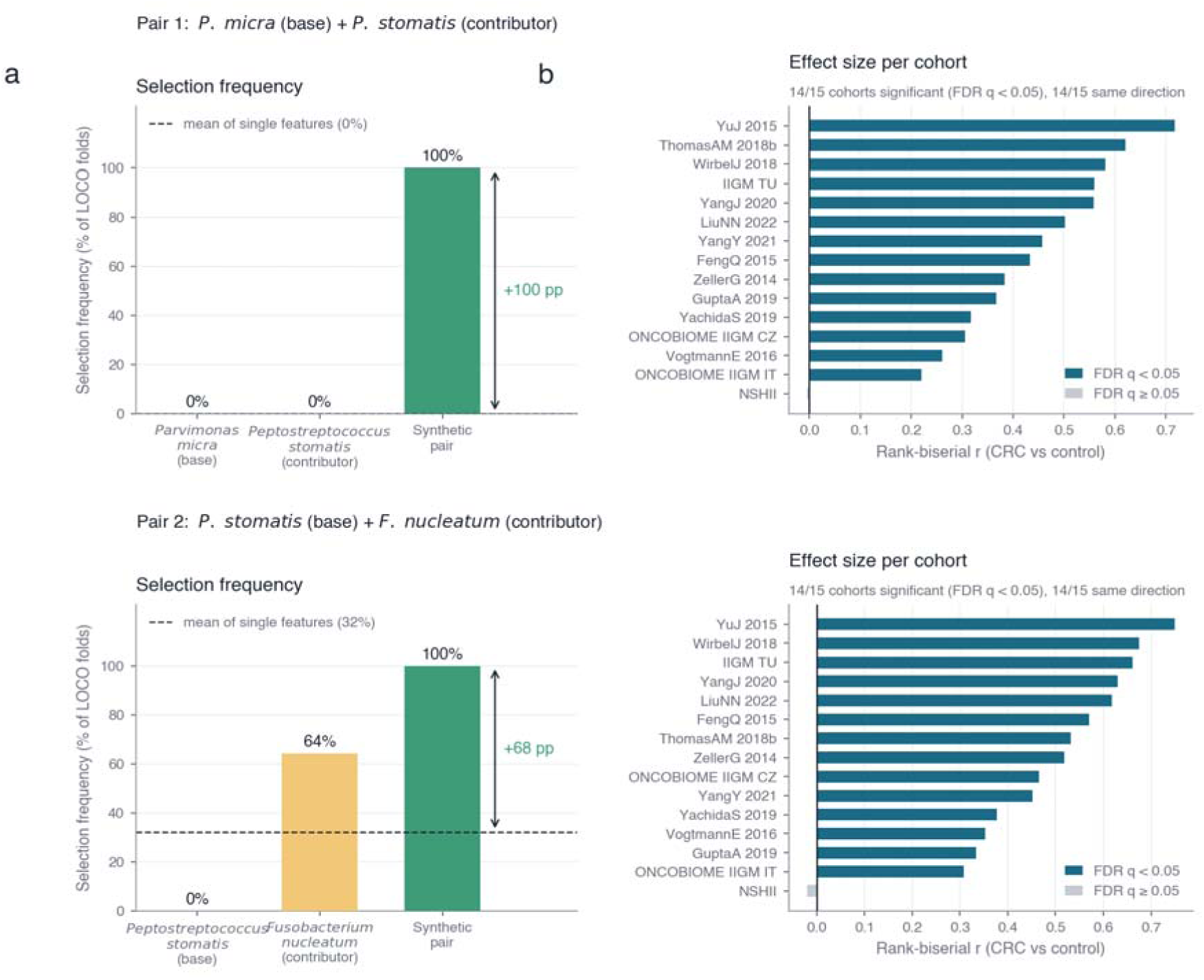
Selection frequency and per-cohort effect sizes of high-frequency synthetic feature pairs. Representative pairs are shown for the two dominant hub species: *Parvimonas micra* (base feature; 0% selection frequency as a raw feature in the combined model) and *Fusobacterium nucleatum* (contributor feature, 64% individual frequency, providing contextual information gain to partners). The two rows show two representative synthetic pairs: pair 1, *Parvimonas micra* (base) + *Peptostreptococcus stomatis* (contributor), and pair 2, *Peptostreptococcus stomatis* (base) + *Fusobacterium nucleatum* (contributor). (a) Frequency of the individual features and of their synthetic pair (green) across LOCO selection folds; the dashed line indicates the mean frequency of the two component features, and the arrow and annotated value show the frequency gain of the pair in percentage points. (b) Post-selection validation using continuous synthetic feature values. Rank-biserial correlations compare synthetic feature values between CRC cases and controls within each cohort (cohorts ordered by effect size); positive values indicate higher values in CRC. In contrast to (a), this analysis takes all sample values into account rather than relying on binary threshold-based splits, providing a complementary, distribution-based measure of effect size; as the same cohorts contributed to pair selection, this analysis supports, but does not independently validate, the selected pairs. Dark teal bars denote statistical significance after multiple testing correction (Benjamini– Hochberg FDR across cohorts within each pair; q < 0.05), grey bars non-significance (q ≥ 0.05).

To assess whether the selected pairs reflected meaningful case-control differences beyond model-based selection frequencies, we performed a post-selection, cohort-level effect size analysis. For each of the 235 stable pairs, we computed rank-biserial correlations between CRC cases and controls in each cohort individually (Figure 2b), using the full distribution of synthetic feature values within each cohort. As the same cohorts contributed to the training data from which the pairs were selected, this analysis provided supporting evidence rather than a fully independent validation. All 235 pairs reached meta-significance across cohorts (sample-size-weighted Stouffer’s method on signed Z-scores; Benjamini–Hochberg FDR across the 235 pairs, q < 0.05), and 225 showed consistent directionality of effect in at least 14 of 15 cohorts (133 in 15 of 15; the remaining 10 in 12-13 of 15). At the individual cohort level, 78.0% of cohort-pair tests were significant after Benjamini–Hochberg correction across cohorts within each pair (q < 0.05), indicating that the selected pair features differ consistently between cases and controls within individual cohorts (Supplementary Table 4).

To check that these differences were not explained by cohort membership, we additionally fitted cohort-adjusted logistic regression models to the pooled samples of the 15 two-class cohorts (log-transformed relative abundances, cohort as a fixed effect; Methods). 235 of the 235 pairs remained informative beyond cohort membership (likelihood-ratio test of the additive model against the cohort-only model, Benjamini–Hochberg q < 0.05), and 123 showed a significant multiplicative interaction term between the two constituents in addition to their additive effects (Supplementary Table 5).

### Pair transformations reveal distinct forms of disease-associated dependence

Because each MDFS-selected pair was encoded using two transformations, we examined which form of dependence the classifier preferentially relied on. The geometric mean (GM) is high when both members are jointly abundant and therefore reflects joint enrichment, whereas the log ratio (LR) reflects the relative dominance of one feature over the other. Joint enrichment is compatible with, but does not demonstrate, mutualism, cross-feeding or a shared niche preference; likewise, a shift in the ratio may, but need not, reflect competition. Providing both transformations to the RF model allowed the algorithm to select the representation that best captured the disease-associated dependence within each pair.

We assessed the predictive utility of these approaches by monitoring their selection frequency within the RF models. While the choice of transformation had negligible impact on many feature pairs, several high-stability variables (selected in ≥80% of folds) exhibited a strict dependence on a specific mathematical form. For instance, the pairs of *P.stomatis* and *R.intestinalis* (93% selection), *D.pneumosintes* and *R.intestinalis* and *P.micra* and *R.intestinalis* were relevant exclusively as log-ratios, with per-cohort effect sizes of the LR encoding far exceeding those of the GM encoding (Figure 3a-c). Conversely, *D. pneumosintes* and *P.stomatis*, *F.nucleatum* and *P.stomatis* and *D.pneumosintes* and *G.morbillorum* emerged solely through the GM transformation (Figure 3d-f), as did the functional pairs of *L-lysine fermentation (P163-PWY)* with the *TCA cycle VI (REDCITCYC)* and other pathways (Figure 3g-i). Some pairs were informative in both forms (71 of the 235 stable pairs, e.g. *A.obesiensis* and *F.nucleatum*). Overall, the predominance of GM-based synergies (173/235 pairs, 74%; GM only 102, LR only 62, both 71) suggested that joint enrichment of specific feature pairs was more consistently associated with CRC status than shifts in their relative balance (Supplementary Table 6).

**Figure 3.**
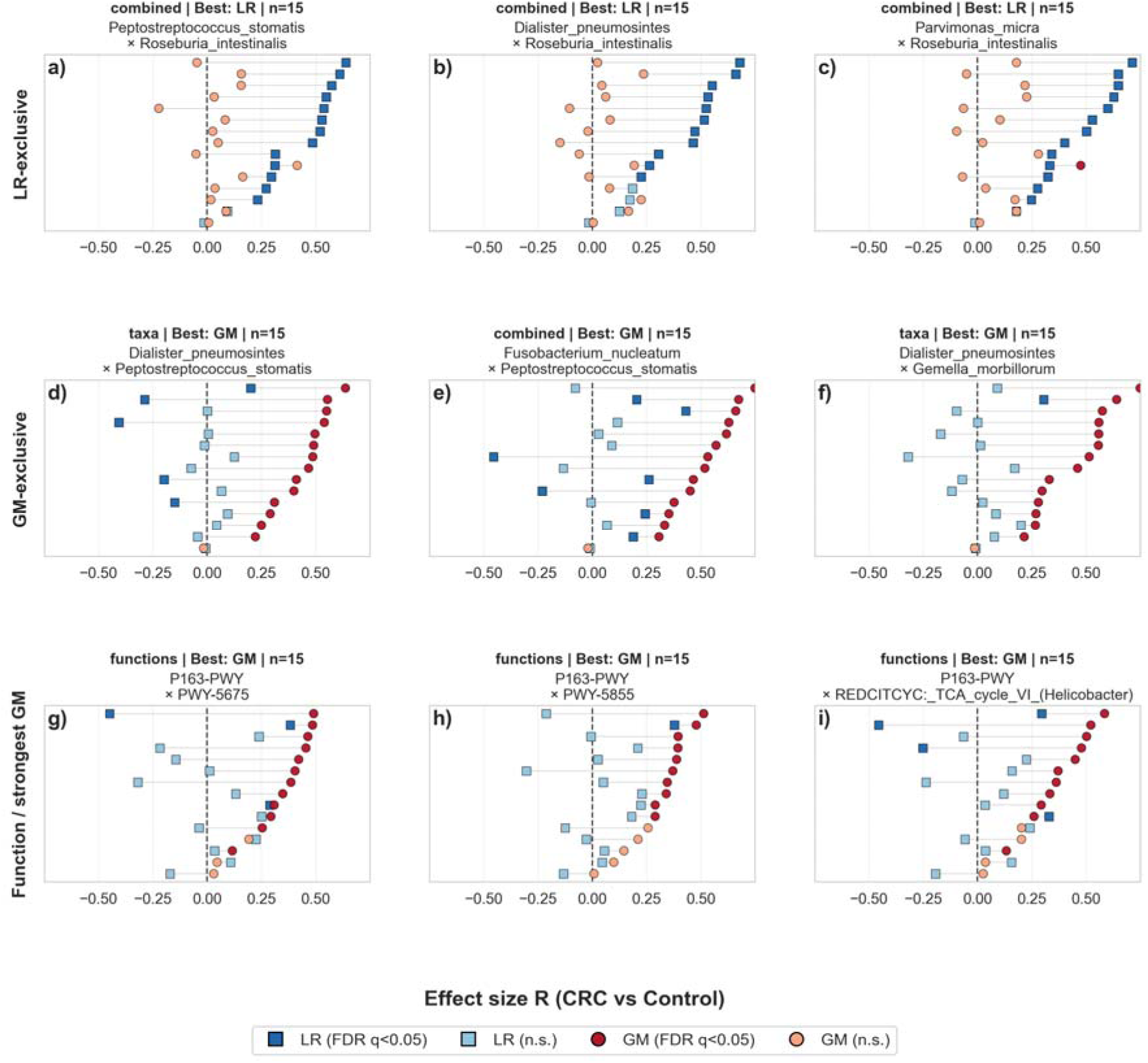
Log-ratio (LR, squares) versus geometric-mean (GM, circles) encodings of nine stable synthetic pairs: a-c) pairs selected exclusively as LR, d-f) pairs selected exclusively as GM, g-i) functional pairs; the pairs with the largest contrast between the two encodings were chosen automatically among the stable pairs. Cohort-specific effect sizes (rank biserial correlation; positive values indicate higher synthetic feature values in CRC relative to controls), with filled dark symbols indicating statistical significance after within-pair multiple testing correction across cohorts (Benjamini–Hochberg FDR; q < 0.05) and light symbols non-significant values. The functional pairs are g) *P163-PWY with PWY-5675 (nitrate reduction V, assimilatory*); h) *PWY-5855 (ubiquinol-7 biosynthesis, early decarboxylation*; and i) *REDCITCYC (TCA cycle VI, Helicobacter)*.

### Broader application identifies synergistic pairs in other conditions, with limited cross-cohort recurrence

Having established that MDFS-2D identifies synergistic feature pairs with genuine predictive value in CRC, we asked whether this synergistic structure was a general property of microbiome-disease associations or specific to colorectal cancer. To explore this, we applied the same framework to a broader set of well-characterized disease cohorts, spanning conditions including inflammatory bowel disease, type 2 diabetes, and liver cirrhosis. Our goal was not to build optimized classifiers for each disease, but rather to determine whether meaningful synergistic pairs - those in which the 2D IG of the base feature substantially exceeds the IG in 1D - emerge consistently across diverse disease contexts.

To test this, we applied our framework to 13 cohorts previously processed through the curatedMetagenomicData set (Table 2 in Methods, (Pasolli et al., 2017)). The pairs were identified using MDFS-2D with the same Bonferroni-corrected pair-level significance threshold and synergy filter as in the CRC discovery pipeline, with 10 MDFS runs per cohort and a pair reported only if it passed in at least 6 runs (Methods). As a result, 8 of the examined 13 cohorts yielded synergistic pairs (80,790 unique pairs in total). Of these, 76,828 (95.1%) exhibited at least a 100% gain in 2D IG over the base constituent’s individual signal in at least one cohort. The remaining five cohorts (HMP_2019_ibdmdb, HMP_2019_t2d, KarlssonFH_2013, LiJ_2017, QinJ_2012) yielded no pair that passed the thresholds in at least 6 of 10 runs, consistent with the conservative Bonferroni correction across ∼1.5 million tested pairs.

The range of synergistic effects observed was broad. Unexpectedly, a large share of pairs was due to a synergy with a strong partner. This was especially pronounced in the QinN_2014 liver cirrhosis dataset, where *C.amalonaticus*, with a negligible individual IG of 0.23, reached a joint IG of 88.2 when paired with *S.anginosus* group species (a 385-fold amplification). As the partner alone carried 81.6, this pair illustrated the directional nature of the gain - the information was largely that of the contributing feature, conditioned on *C.amalonaticus*. In other cases, however, the pairwise search found features that a univariate search could not detect. In 3,808 pairs, one constituent carried no detectable individual signal whatsoever - yet the pair yielded substantial joint relevance when the silent feature was combined with an informative partner. We term these one-sided "emergent" synergies. Such emergent pairs were observed in two cohorts: HallAB_2017 and MetaCardis_2020_a. The most dramatic gains involved features with weak but non-zero individual signal that underwent substantial amplification in combination. In the HallAB_2017 IBD cohort, *E.ramulus* carried a modest individual IG of 0.58. However, when paired with *R.torques* (individual IG 26.7) as a contributing feature, the 2D interaction added 54.3 to reach a joint IG of 54.9 - a 94-fold amplification of the base signal and twice the information of the partner alone. Synergistic signal extended beyond species-species interactions: in the MetaCardis_2020_a cardiometabolic cohort, the top-scoring pair combined *superpathway of L-aspartate and L-asparagine biosynthesis (ASPASN-PWY)* with *adenine and adenosine salvage III (PWY- 6609)* (joint IG = 61.0). Together, these examples illustrate that synergistic signal was not confined to a single disease or feature type but emerged broadly and with considerable magnitude across diverse microbiome-disease contexts.

To understand how broadly and strongly synergistic signal manifested across disease contexts, we examined the distribution of synergistic IG gains across the eight cohorts with significant 2D results. The number of synergistic pairs with a >=50% IG gain varied considerably, from a single pair in ZhuF_2020 and 2 in NagySzakalD_2017 to 55,850 in QinN_2014 and 12,049 in HallAB_2017 (Figure 4b; see Table 2 counts for all retained pairs, e.g. 7,585 in MetaCardis_2020_a, of which 3,805 reach a 50% gain). This variation mainly reflected differences in sample size, disease effect size, and the stringency of the Bonferroni correction across cohorts.

**Figure 4.**
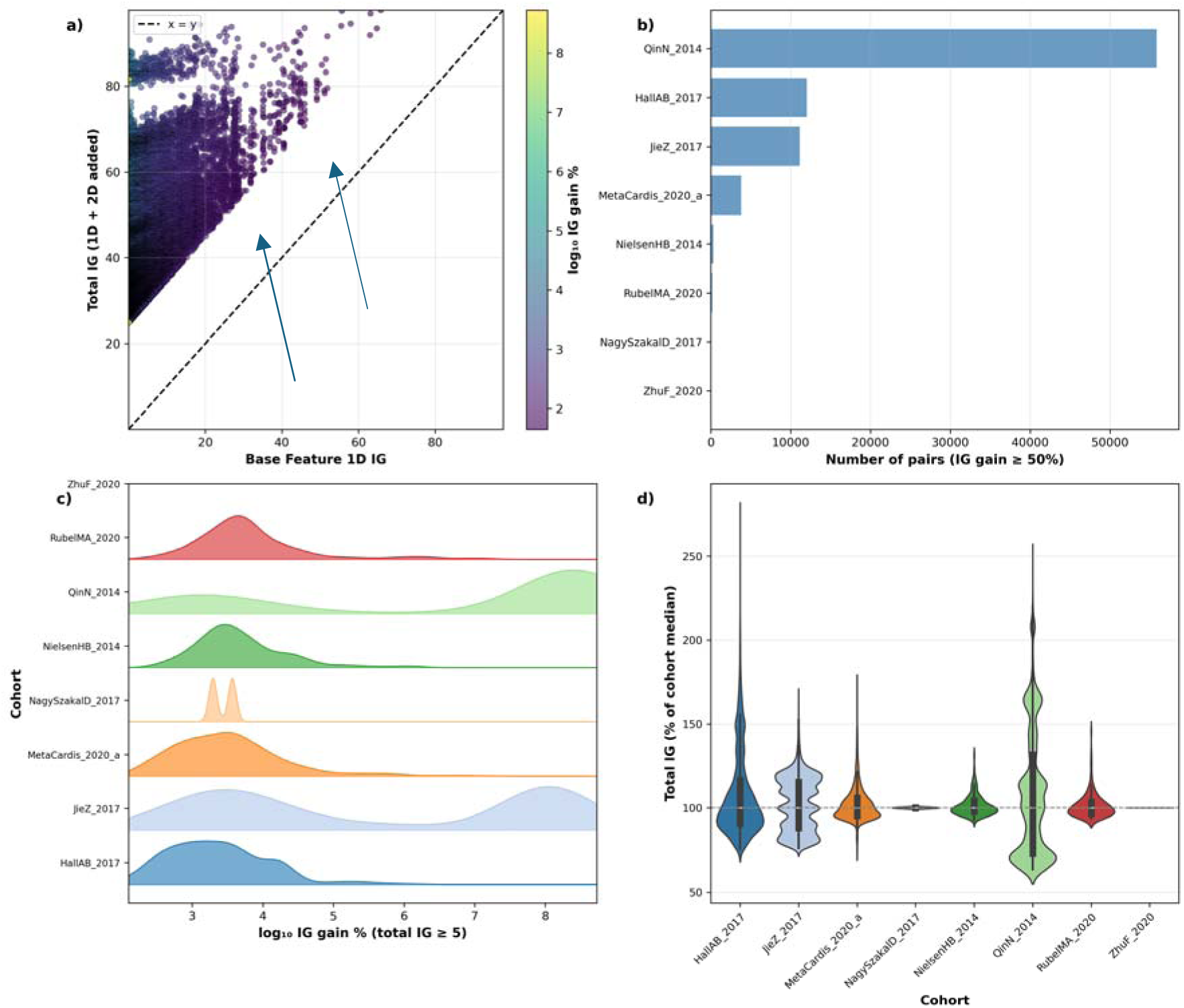
2D information gain and pair-level synergy across validation cohorts. a) Joint IG of each synergistic pair (1D IG of the base feature + IG added by the contributing feature) versus the 1D IG of the base feature; one point per unordered pair and cohort (orientation with the larger joint IG), coloured by the relative gain; points above the dashed line (x = y) show positive 2D gain. b) Number of unordered pairs per cohort with at least 50% IG gain. (c) Ridge plot of IG %, restricted to pairs with total_IG >= 5. (d) Violin plots of total_IG as % of cohort median. Colours match (c).

Among cohorts with substantial pair counts, the QinN_2014 liver cirrhosis and JieZ_2017 ACVD datasets stood out for both the number and strength of synergistic interactions, with nearly all pairs (99.7% and 100.0%, respectively) exhibiting at least a doubling of the base feature’s IG (Figure 4c). At the other end of the spectrum, MetaCardis_2020_a - which encompassed multiple cardiometabolic conditions - showed the most modest proportion of high-gain pairs (50.2% with >=100% gain), potentially reflecting a more heterogeneous patient population. No pair recurred in more than three cohorts: 6,232 pairs were found in at least two cohorts and 102 in three (68 of them in HallAB_2017, JieZ_2017 and QinN_2014, and 28 in JieZ_2017, MetaCardis_2020_a and QinN_2014). 89% of the latter involved an oral *Streptococcus* species (*S.vestibularis*, *S.mitis*, *S.anginosus* group or *S.oralis*) paired with a gut commensal or a pathway, and 9 of the remaining 11 pair a gut species or pathway with *adenine and adenosine salvage III (PWY-6609)*.

Finally, we investigated whether synergistic pairings were conserved across independent cohorts sharing the same clinical diagnosis, which would point toward candidate disease-specific synergistic markers (Supplementary Table 7). For this we decided to include pairs which were matched irrespective of which member MDFS reported as the base feature, because the same pair could pass the per-direction thresholds in opposite orientations in different cohorts. Ultimately, cross-cohort comparison was possible only for IBD (HallAB_2017, NielsenHB_2014). We identified 14 conserved IBD pairs detected in both independent cohorts, comprising 10 taxa-taxa and 4 taxa-function pairs, all of which involved *B.hydrogenotrophica*. The strongest conserved IBD synergies were *A.putredinis* with *B.hydrogenotrophica* (mean joint IG 39.5 across both cohorts), *B.hydrogenotrophica* with *phosphopantothenate biosynthesis I (PANTO-PWY)* (34.0) and *B.intestinalis* with *B.hydrogenotrophica* (34.0). Notably, the orientation differed between the cohorts for every one of these pairs: *B.hydrogenotrophica* carried no individual signal in HallAB_2017 and gained all its information from the partner, whereas in NielsenHB_2014 it was the partner that was conditioned on *B.hydrogenotrophica*. No cross-cohort comparison was possible for type 2 diabetes, as none of the three T2D/IGT cohorts analysed (HMP_2019_t2d, KarlssonFH_2013, QinJ_2012) yielded pairs that passed the thresholds in at least 6 of 10 runs.

## Discussion

In this study, we introduced a framework for identifying synergistic microbial feature pairs using the Multidimensional Feature Selection (MDFS) algorithm. Our work was motivated by a fundamental limitation of conventional approaches: the reliance on univariate analysis that evaluates metagenomic markers in isolation, overlooking the combinatorial signals that arise from microbial co-occurrence. Validating on a well-characterized CRC meta-analysis dataset (Piccinno et al., 2025) - chosen for its rigorous published benchmark and clinical relevance as a target for non-invasive microbiome-based diagnostics - our framework selected feature pairs on which synthetic-only random forest models reached performance comparable to the benchmark (taxonomy AUC = 0.84 vs 0.85; functions-only AUC = 0.67). The goal of this exercise was not to outperform the best models, but rather to show that synergy-based feature selection could add a layer of biological interpretability without compromising performance.

Why therefore use a synergy-oriented feature selection framework when predictive performance is unchanged? High-performing classifiers such as random forests implicitly exploit feature interactions but do not report them in a form that can be inspected or tested. Model-based attribution methods, such as interaction terms in regression models or SHAP interaction values, explain a fitted model; the pairs they report would be expected to be among the pairs scanned exhaustively here. Our framework makes these dependencies explicit: it returns the specific pair, the partner providing the additional information, and the transformation (co-abundance or ratio) through which the pair is informative, together with its stability across held-out cohorts. Our aim was all-relevant discovery rather than a minimal predictive subset: a pair was reported because it was statistically relevant and showed how disease-associated information was distributed among features. Such outputs are hypothesis-generating rather than confirmatory, and - if validated - they naturally translate into compact diagnostic designs. This includes targeted assays quantifying a small number of taxon pairs or ratios, which are less sensitive to variation in total microbial load than absolute single-taxon abundances.

Our framework consistently identified synergistic pairs in which the 2D IG of the base feature substantially exceeded its individual IG in isolation. Several of these combinations were also highly prevalent, selected as relevant independently across multiple CRC cohorts. Particularly striking was the dominance of *F. nucleatum* among high-IG synergistic pairs, where it appeared consistently partnered with oral cavity anaerobes - *P. stomatis*, *P. micra*, and *D. pneumosintes* - all established members of the subgingival niche. These findings align with the increasingly recognized oral microbiota signature in CRC (Chen et al., 2025) . Oral microbiota may contribute to CRC pathogenesis through multiple mechanisms: direct translocation via daily swallowing, mucosal barrier breach enabling systemic bacteremia, and immune modulation through colitogenic pathobionts promoting pro-inflammatory Th17 responses (Sears and Garrett, 2014; Tanoue et al., 2019). Our observational data, however, cannot distinguish whether these taxa contribute causally to tumorigenesis or become enriched secondarily in the tumour-associated environment. Many of these taxa are established CRC markers in their own right (Piccinno et al., 2025).

Strikingly, the high-stability pairs were not a collection of independent pairings but a densely interconnected network. *F.nucleatum* and *P.micra* appeared repeatedly across many combinations (125 and 48 stable pairs across the three feature sets, respectively) - both as contributing features and as base-feature hubs - each co-occurring with multiple other oral anaerobes in overlapping configurations. These pairs may represent cross-sections of a single higher-order oral community signature, in which the meaningful co-occurrence of multiple oral species, rather than any individual taxon, was associated with CRC (Dejea et al., 2014; Flynn et al., 2016) . This hypothesis was consistent with evidence that oral bacteria detected in CRC tissue co-occur in spatially organized biofilm structures, co-occur during tumour progression and are commonly identified in metastasis (Bullman et al., 2017) , as well as exhibit cross-species metabolic dependencies that sustain the community in the anaerobic colonic microenvironment (Hajishengallis and Lamont, 2012) .

To characterize the form of the disease-informative dependence identified within each synergistic pair, we encoded pairs using two complementary transformations. The geometric mean (GM) is high when both members are jointly abundant, whereas the log-ratio (LR) reflects the relative dominance of one feature over the other. These transformations described the structure of the dependence - joint enrichment versus relative balance - but did not by themselves identify the underlying ecological process. Joint enrichment could arise from mutualism or cross-feeding, but equally from a shared environmental preference or a common response to the host, while a shift in ratio did not necessarily imply competition. The predominance of GM-based features among stable pairs indicated that, in this dataset, joint enrichment of feature pairs was more informative of CRC status than changes in their relative balance. Whether specific GM- or LR-encoded pairs reflected actual biological relationships between their members was an important direction for future work and would require validation using independent approaches such as co-culture experiments, metabolic modelling, or longitudinal and absolute-abundance data.

The emergent synergies identified in the broader cohort analysis show what the 2D framework can detect beyond 1D analysis: pairs in which one constituent showed no individual signal became informative when conditioned on an informative partner. In these pairs the total IG largely reflects the informative partner, with the silent member acting as a context variable (its presence or absence) rather than contributing signal of its own. Such patterns are compatible with ecological dependencies, for example one species modulating local conditions or supplying metabolic substrates for another (Fischbach and Sonnenburg, 2011), but they are equally compatible with shared responses to host factors. Because cohorts were not tested against a null model, these pairs should be treated as hypotheses, and distinguishing between the explanations requires experimental validation.”

Together, our findings exposed a systematic blind spot in standard microbiome analyses: univariate approaches do not report the interaction-level signal that accounts for a substantial share of disease-relevant microbiome structure, and in the CRC data the pairwise scan recovered almost every feature that univariate tests selected while adding features that none of them did (Supplementary Figure 1). The recurrence of pairs across 3 cohorts and of emergent synergies in 2 cohorts suggested that these were not merely cohort-specific artefacts and supported the existence of a layer of disease-relevant microbiome structure that pairwise analysis resolved.

### Limitations

Several limitations of this study should be noted. First, in the CRC benchmark, synergy-derived features did not improve classification performance over raw features. Instead, their value was in making informative feature dependencies explicit rather than in predictive gain; moreover. Functional pathway profiles alone showed limited discriminative power in this benchmark (AUC = 0.67). Second, although the framework identified meaningful, disease-informative co-occurrence patterns rather than co-occurrence per se, these remain statistical dependencies: they provide no information on causality or on the nature of the underlying biological interactions, and the ecological interpretations discussed here are hypotheses that require experimental validation. Third, cross-cohort conservation could be assessed within a single diagnosis only for IBD, and the overlap of synergistic pairs across cohorts and diseases was limited. The recurring pairs reported here are therefore candidate markers that require validation in independent cohorts before any diagnostic application. Fourth, the per-cohort effect-size analysis was performed on cohorts which had also contributed to pair selection and was therefore a post-selection characterization. Classification performance, in contrast, was always evaluated on held-out cohorts, and confirmation of the stable pairs in a cohort not used for selection would strengthen them further. Fifth, the analysis operated on relative abundances. MDFS discretises each feature independently and evaluates the information carried by the resulting bins, so it does not assume any particular scale. The compositional nature of the data nevertheless meant that a dependence between two features could partly reflect the shared constraint of the simplex, and the log-ratio encoding was the only one of the two transformations that was invariant to it. Most taxonomic features were zero-inflated (76% had a median abundance of zero), so the binning largely reflected presence versus absence, and no taxon systematically dominated the samples (the most abundant taxon had a median of 17.5% of a sample). Sixth, the pair search was exhaustive only at the level of pairs - higher-order combinations, which MDFS can in principle evaluate, were not explored. Seventh, in each LOCO fold the feature space was restricted to the features present in every training cohort (Methods), which favoured prevalent taxa and pathways and made the feature space depend on the training cohorts. Finally, feature selection in the functional feature space was markedly less stable across MDFS seeds than in the taxonomic space. A negative control delimited what the pair search detected: with permuted labels the pipeline returned at most 1 pair (real data: 31,751–183,176 pairs per feature space), so the retained pairs were not false positives of the significance test. The discretisation used by MDFS (two bins per feature at a random cut point) was a modelling choice; on one LOCO fold, widening or halving the range of admissible cut points changed the set of synergistic pairs by about as much as a change of random seed (Jaccard of all passing pairs with the default 0.96 and 0.90, versus 0.78 between seeds), whereas using three bins per feature changed it more substantially (Jaccard 0.66). The per-run top 2% of pairs, from which the consensus is built, was far less stable in a single run (Jaccard 0.14 between seeds, 0.17 for the halved range and 0.26 for three bins; Supplementary Table 8), which is why repeated runs and a consensus rule are part of the framework.

### Conclusion

Collectively, our results establish pairwise feature synergy as a reproducible and biologically informative axis of microbiome-disease association that standard univariate approaches do not resolve. By treating the feature pair rather than the individual feature as the fundamental unit of analysis, the MDFS-based framework recovers a layer of disease-relevant signal - from hub taxa whose signal is captured preferentially at the pair level to emergent synergies in which one constituent carries no detectable individual information. The recurrence of these signals across feature types and, for a small set of pairs, across independent cohorts suggests that meaningful microbial co-occurrence structure represents a genuine and underexplored dimension. As microbiome research moves from cataloguing associations toward understanding the interaction networks that underlie them, frameworks capable of capturing combinatorial rather than marginal effects will be essential - both for identifying interpretable biomarker candidates and for designing interventions that target the community rather than the individual taxon.

## Materials and Methods

### Data preparation

Metagenomic data was obtained from a recently published CRC meta-analysis benchmark (Beghini et al., 2021; Piccinno et al., 2025) . Briefly, the dataset comprised human gut metagenomes from 17 independent cohorts of healthy individuals and CRC patients, spanning multiple countries across Europe, Asia, and North America. Taxonomic profiles were generated using MetaPhlAn 4 (v4.0.0) (Blanco-Míguez et al., 2023) and functional profiles using HUMAnN 3.6, applied consistently across all cohorts (Beghini et al., 2021) . Raw metagenomes were preprocessed to remove low-quality reads, short fragments, and host-derived sequences prior to profiling. For our analysis, MetaPhlAn species-level (*s* ) relative abundances constituted the taxonomic feature set; genome-level SGB bins (*t* ) were excluded, and species-level labels were used for consistency with the benchmark study (Piccinno et al., 2025). Pathway abundances were converted to within-sample relative abundances; for the combined feature space, species-level taxa and pathway features were concatenated into a single matrix and re-normalized to relative abundance per sample so that the taxonomic and functional blocks each contributed equally (summing to 0.5) to every sample. No additional prevalence or abundance filter was applied within cohorts. In each LOCO fold, however, the feature matrix was restricted to the features present in every training cohort, so that MDFS and the random forest operated on a common feature space: 473–484 species (of 821–1,957 reported per cohort), 445–461 pathways and 918–930 combined features per fold, and 682, 461 and 1,143 in the fold in which the cohort with the fewest reported species was held out. Features of the held-out cohort outside this space were ignored and features of the space absent from the held-out cohort were set to zero. Feature relevance within this space was assessed directly by MDFS. A full description of cohort demographics, sequencing procedures, and preprocessing pipelines is provided in the original publication (Piccinno et al., 2025) . The 17 cohorts and their numbers of CRC and control samples are listed in Table 1.

**Table 1.** CRC cohorts of the discovery arm (the 17 cohorts of the CRC meta-analysis benchmark), with the numbers of CRC and control samples used in the analysis. AtezoTRIBE, COLOBIOME, IIGM_CZ, IIGM_IT, NSHII and IIGM_TU are in-house cohorts of this study. †Cohorts never held out as test sets, which entered the training set of every LOCO fold (AtezoTRIBE and COLOBIOME contain CRC samples only; NSHII has 14 CRC samples, below the minimum of 15).

| Cohort | CRC (n) | Control (n) | References |
| --- | --- | --- | --- |
| FengQ_2015 | 46 | 61 | (Feng et al., 2015) |
| GuptaA_2019 | 30 | 30 | (Gupta et al., 2019) |
| LiuNN_2022 | 85 | 78 | (Liu et al., 2022) |
| ThomasAM_2018b | 28 | 32 | (Thomas et al., 2019) |
| VogtmannE_2016 | 52 | 58 | (Vogtmann et al., 2016) |
| WirbelJ_2018 | 60 | 65 | (Wirbel et al., 2019) |
| YachidaS_2019 | 258 | 290 | (Yachida et al., 2019) |
| YangJ_2020 | 95 | 98 | (Yang et al., 2020) |
| YangY_2021 | 100 | 100 | (Yang et al., 2021) |
| YuJ_2015 | 54 | 74 | (Yu et al., 2017) |
| ZellerG_2014 | 53 | 61 | (Zeller et al., 2014) |
| AtezoTRIBE† | 163 | 0 | (Piccinno et al., 2025) |
| COLOBIOME† | 201 | 0 | (Piccinno et al., 2025) |
| IIGM_CZ | 46 | 59 | (Piccinno et al., 2025) |
| IIGM_IT | 59 | 86 | (Piccinno et al., 2025) |
| NSHII† | 14 | 448 | (Piccinno et al., 2025) |
| IIGM_TU | 39 | 18 | (Piccinno et al., 2025) |

**Table 2.**
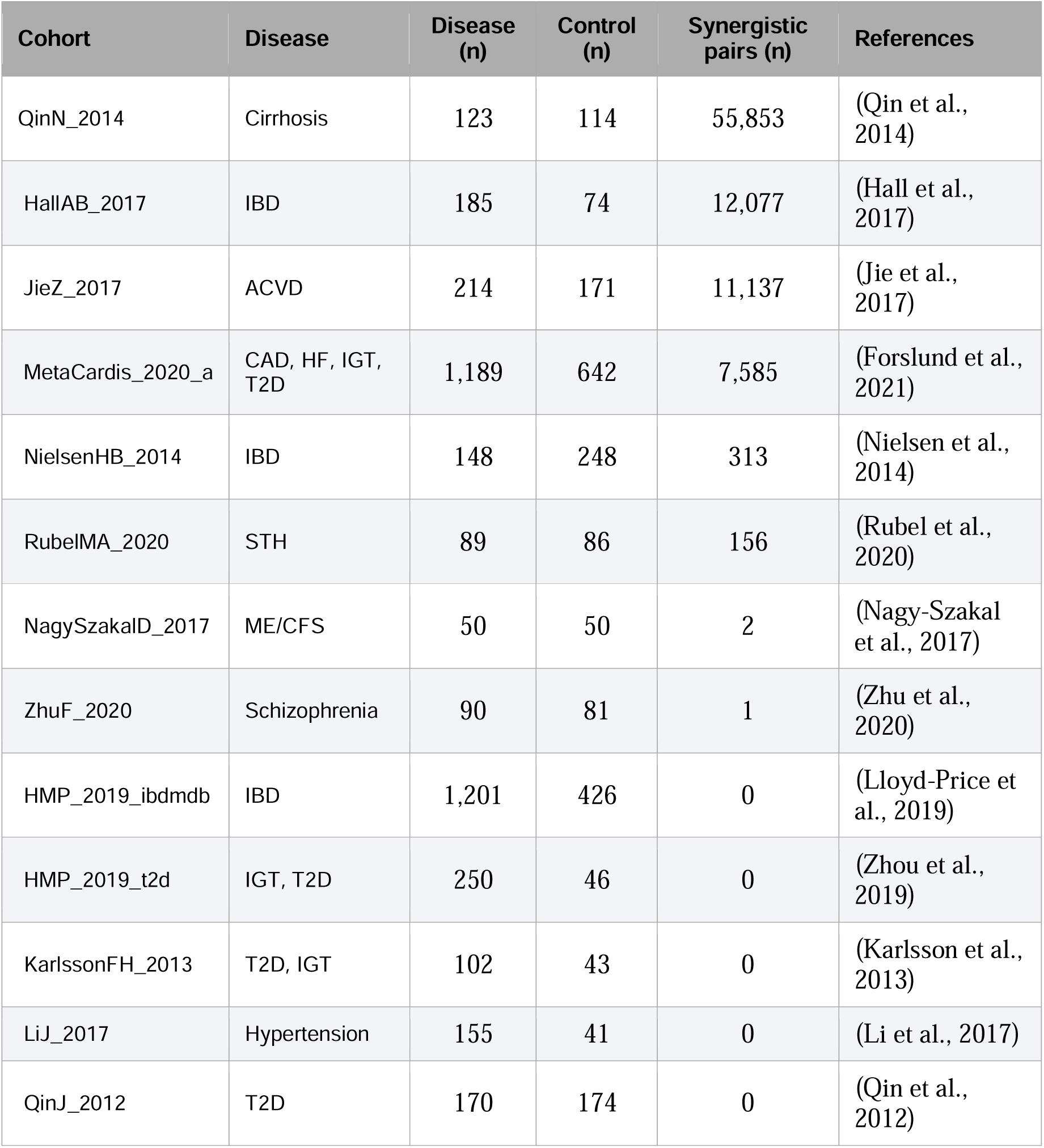
Non-CRC cohorts analysed in the broader application, ordered by the number of synergistic pairs (unique pairs that passed the pair-level significance and synergy thresholds in at least six of ten MDFS runs). Disease (n) and Control (n) give the numbers of samples.

For the broader synergy validation analysis on non-CRC cohorts, the curatedMetagenomicData Bioconductor package (Pasolli et al., 2017), in the most recent version available at the time of analysis (summer of 2026), was queried for non-CRC cohorts containing both disease and control samples and at least ten samples. Twenty such cohorts were identified; taxonomic and functional profiles were available in the downloaded release for 13 of them, which were analysed (Table 2). The remaining seven (KosticAD_2015, LiJ_2014, RaymondF_2016, SankaranarayananK_2015, VatanenT_2016, VincentC_2016, YeZ_2018) had no profiles in the download and could not be analysed.(Pasolli et al., 2017) All profiles in *curatedMetagenomicData* were generated uniformly using MetaPhlAn3 (taxonomic abundances) and HUMAnN3 (metabolic pathway abundances) (Beghini et al., 2021) . The cohorts spanned a broad range of conditions as presented in Table 2.

### MDFS feature selection and information gain

Feature selection was performed with the R package MDFS (v1.5.5) (Piliszek et al., 2019) . MDFS was run in both 1D and 2D modes: MDFS(X, y, dimensions = 1) identified individually significant features (Holm-corrected α = 0.05), returning per-feature information gain (IG) values directly via the test statistic. MDFS(X, y, dimensions = 2) identified features participating in significant pairwise interactions. For the 2D-significant variables, feature pairs were enumerated using ComputeInterestingTuples(X, y, dimensions = 2, interesting.vars = sig_2d_vars, I.lower = ig_1d, ig.thr = ig_thr_pair), where I.lower was set to the vector of 1D IGs so that the returned IG represents the additional 2D information gain (IG_2D_added) contributed by the partner feature beyond the base feature’s marginal 1D IG. IG values are reported as MDFS computes them, in bits summed over the samples of the data set (the chi-squared statistic divided by 2 ln 2, i.e. N times the mutual information). In simple terms, they scale with the sample size and thresholds are specific to a dataset. Pair-level significance was assessed using a chi-squared test (df = 2 under the default MDFS discretisation), with a Bonferroni correction over all n(n−1)/2 possible feature pairs at α = 0.05, yielding the IG threshold ig_thr_pair = qchisq(1 − 0.05/n_pairs, df = 2) / (2·log 2). As an additional selection step, which is not part of the definition of synergy, a pair direction was retained only if its total IG also exceeded the 1D IG of the contributing feature (total_IG > max(base_IG_1D, contributing_IG_1D)). This removed directions in which the gain of a weak base feature merely reflected the individual signal of a strong partner. MDFS discretises every feature into two bins (divisions = 1) at a cut point drawn at random within the range recommended by the package for two-dimensional analyses (GetRange(n, dimensions = 2)). At range = 0 this cut would coincide with the feature’s median, whereas the wide recommended range draws it at random from a broad window around the median, so that for the many taxa whose median abundance is zero the split reduces to presence versus absence.

The same seed and cut range were used for the 1D, 2D and pair-level computations, so that each feature is binned identically in all steps and the joint IG of a pair decomposes exactly into the 1D IG of the base feature plus the IG added by the contributing feature. The sensitivity of the pair selection to the discretisation (cut range and number of bins) is reported in Supplementary Table 8, and a negative control (permuted labels) in Supplementary Table 9. One MDFS run on the pooled training set of a LOCO fold (2,834 samples) took 3 s for 473 taxonomic features (111,628 candidate pairs), 2 s for 445 functional features and 9 s for the 918 combined features (420,903 candidate pairs), with a peak memory of 0.4 GB, on a Apple M4 Pro (14 cores); the complete LOCO analysis (14 folds × 10 runs × 3 feature spaces plus random-forest training) took a few hours on a desktop computer.

Comparison with other feature-selection methods. On the pooled training set of each LOCO fold and for each feature space, the set of relevant features was determined with six selectors: MDFS-1D and MDFS-2D (features relevant at Holm-corrected α = 0.05 in at least six of the ten runs with seeds 42–51, with the discretisation of the main pipeline), Welch’s t-test and the Mann–Whitney U test on log10-transformed abundances (Holm-corrected α = 0.05), L1- penalised logistic regression (LASSO; standardised log10 abundances, class-weighted, liblinear solver with at most 300 iterations; features with a non-zero coefficient) and ReliefF (k = 10 nearest hits and misses, Manhattan distance on min–max-scaled log10 abundances. As ReliefF provides no significance test, the top-ranked features were taken, as many as MDFS-2D selected in the same fold), and, as a size-matched univariate reference, the N features with the highest mean 1D IG over the ten MDFS runs, N being the number of MDFS-2D features of the fold (’1D top-N’). For ReliefF and 1D top-N the two coverages are therefore identical by construction and measure overlap. The coverage of method A by method B is the share of the features selected by A that B also selected, averaged over the 14 folds (Supplementary Figure 1). This comparison concerns individual features.

### Leave-one-cohort-out (LOCO) validation

Evaluation was done in a leave-one-cohort-out (LOCO) framework and the dataset filtering, as well as the two-step Random Forest (RF) training (to select relevant variables and to make final classification) followed that introduced in another study (Piccinno et al., 2025) . Of the 17 CRC cohorts, 14 were eligible to serve as held-out test sets (requiring ≥15 CRC and ≥15 control samples). The remaining three cohorts were never held out and were included in the training set of every fold: the two single-class ONCOBIOME cohorts (AtezoTRIBE and COLOBIOME), which contain CRC samples only, and NSHII (14 CRC samples), which was designated as training-only. Eligibility therefore determined which cohorts could be held out, but did not restrict training. In each fold, MDFS and all subsequent feature selection steps, including the internal selection of k, were applied exclusively to the training cohorts, and the held-out cohort was used only for evaluation. The per-cohort effect size analysis of stable pairs was performed post-selection on the 15 cohorts containing both classes.

For each LOCO fold, MDFS was run ten times with independent random seeds (42-51) to capture the variability of the discretisation-based pair selection. Within each run, the pairs passing the significance and synergy filters were ranked by their total IG (using the maximum over the two orientations as the score) and the top 2% were retained (about 500-1,200 of 25,000-60,000 pairs for the taxonomic feature space). A pair entered the consensus set of the fold if it was among the top 2% in at least six of the ten runs. The choice of ten runs, the 2% cutoff and the threshold of six was based on a seed-stability analysis in which consensus sets built from two disjoint blocks of ten seeds (42-51 and 52-61) were compared for every fold: with these settings the two consensus sets agreed with a mean Jaccard index of 0.95 (taxa), 0.98 (combined) and 0.33 (functions), compared with 0.47, 0.83 and 0.13 for two runs with both runs required (Supplementary Table 1). The consensus set was used for synthetic feature generation and downstream evaluation.

From each consensus MDFS pair (f1, f2), two synthetic features were computed:

- Log-ratio (LR): LR_{f1} {f2} = log((X[f1] + ε) / (X[f2] + ε))
- Geometric mean (GM): GM_{f1} {f2} = sqrt((X[f1] + ε) × (X[f2] + ε))

A pseudocount ε = 1×10 was added to avoid division by zero and undefined logarithms. Synthetic features were generated separately on training and test data, with test data aligned to training features; missing features were filled with 0.

RF classifiers were trained in a similar manner to an existing benchmark (Piccinno et al., 2025) , using scikit-learn with: n_estimators = 1000, max_features = ’sqrt’, min_samples_leaf = 1, class_weight = ’balanced’, and random_state = 42. The number of features k was chosen by internal validation as follows: i) training data was split into 75% train / 25% validation (stratified, random_state=42); ii) a ranking RF was fit on the full feature set and features were ordered by Gini importance; iii) for k ∈ {50, 100, 150, 200, 250, 300, 350, 400, 450, 500}, RF models were fit using the top k features and evaluated on the validation set; iv) the k with the highest validation AUC was selected; and v) a final RF was trained on the full training set using the top k features and evaluated on the held-out cohort.

Three feature configurations were evaluated within the RF framework: (i) feature-only models, trained exclusively on raw taxonomic or functional features; (ii) synthetic-only models, trained on the computed LR and GM transformations of consensus MDFS pairs; the number of synthetic features is twice the number of consensus pairs of the fold (median per fold: see Supplementary Table 1); and (iii) combined models, in which synthetic features were appended to the raw feature matrix prior to the RF training procedure. In combined models, both raw and synthetic features were subject to the same Gini-importance ranking and internal k-selection procedure.

Model performance was assessed on each held-out cohort using the area under the receiver operating characteristic curve (AUC-ROC) and accuracy. Summary performance values (mean and sample-size-weighted mean across test cohorts) were reported for each feature configuration and data type.

### Feature frequency analysis

For each synthetic feature, selection frequency was defined as the proportion of LOCO folds in which it appeared in the final RF model. To focus on robust associations, only synthetic features with a selection frequency ≥ 80% across folds were retained for downstream analysis. For each high-stability synthetic feature, a frequency gain was computed as the difference between its fold-selection frequency and the mean fold-selection frequency of its two constituent raw features. Pairs were retained only if (i) the frequency gain was positive and (ii) the synthetic feature’s selection frequency exceeded that of both individual component features. This deterministic filter ensures that the pair is more consistently selected by the classifier than either constituent alone, confirming that the interaction - rather than either marginal feature - drives model reliance.

### Post-selection effect size analysis

Cohort-level effect sizes for synthetic features were estimated using the rank-biserial correlation, derived from a two-sided Mann–Whitney U statistic comparing synthetic feature values between cases and controls within each cohort. Positive values indicated higher synthetic feature values in CRC cases relative to controls. For each pair, both log-ratio and geometric-mean transformations were evaluated; the transformation available for the most cohorts was selected. P-values across cohorts were corrected for multiple testing within each pair using the Benjamini–Hochberg procedure; cohorts with FDR-adjusted q < 0.05 were considered statistically significant. This approach constitutes the analysis underlying Figure 2b and provides a continuous, distribution-based complement to the RF-based selection frequency analysis in Figure 2a: whereas the RF framework uses binary feature splits to assign group membership, the rank-biserial correlation uses all sample values and is therefore sensitive to the full range of abundance differences between cases and controls. This analysis was performed post-selection on the 15 cohorts containing both classes, all of which contributed to the training data used for pair selection. It therefore provided supporting, post-selection evidence for the selected pairs rather than a fully independent validation. To summarize evidence across cohorts, signed Z-scores from the per-cohort tests were combined using Stouffer’s method weighted by the square root of cohort sample size. The resulting two-sided meta p-values were corrected across pairs using the Benjamini–Hochberg procedure. To assess whether the stable pairs remain informative after adjustment for cohort membership, logistic regression models were fitted to the pooled samples of the 15 two-class cohorts for every stable pair, with log10-transformed relative abundances (pseudocount 10^-6^) of the two constituents and cohort as a fixed effect. Likelihood-ratio tests compared the additive model with the cohort-only model and the model with a multiplicative interaction term with the additive model, with Benjamini–Hochberg correction across pairs (Supplementary Table 5). Spearman correlations between each stable synthetic feature and its constituents were computed on the same pooled data. The 1D significance of individual features was taken from the Holm-adjusted p-values returned by MDFS in 1D mode for every fold and run. As a sensitivity analysis of the training-set composition, the complete LOCO pipeline was repeated with the three cohorts that were never held out (AtezoTRIBE, COLOBIOME, NSHII) removed from every training set (Supplementary Table 3).

### Application to additional disease cohorts

To evaluate the generalizability of the synergy-detection framework, MDFS was applied independently to each non-CRC disease cohort without a LOCO framework, using the full cohort as input. MDFS was run in both 1D and 2D modes ten times with independent random seeds (12–21), applying the same statistical filters as in the discovery arm: Bonferroni-corrected chi-squared pair-level significance and the synergy filter (total_IG > max(base_IG_1D, contributing_IG_1D)). A (base, contributing) direction was retained if it passed both filters in at least six of the ten runs, and its IG values were averaged over the runs in which it passed. Because MDFS reports a pair in each direction that passes the thresholds, with different base and added IG but the same joint IG, all pair counts refer to unordered pairs, represented by the direction with the larger joint IG. For each pair, the synergy gain percentage was computed as ig_gain_pct = IG_2D_added / base_IG_1D × 100, representing how much additional information the contributing feature provided relative to the base feature’s individual IG. Pairs with ig_gain_pct ≥ 50% were retained for downstream analysis. Pairs in which at least one feature had zero 1D IG (base_IG_1D = 0 or contributing_IG_1D = 0) yet exhibited positive 2D IG were designated "emergent" synergies. This term was defined as interactions in which one feature carried no detectable marginal signal and became informative only in the presence of its partner. To identify conserved synergistic markers, unordered pairs observed in ≥ 2 independent cohorts sharing the same disease label were retained.

## Data Availability

The CRC cohort was identical to the one in the recently published CRC meta-analysis benchmark (Beghini et al., 2021; Piccinno et al., 2025) and is listed in Table 1; the other datasets included in this study are listed in Table 2. All datasets were retrieved from the *curatedMetagenomicData* Bioconductor package (Pasolli et al., 2017) , using the most recent version available at the time of analysis.

All code, with one folder per manuscript section and the files behind every figure and table, is in the *manuscript* branch of https://github.com/Kizielins/MDFS_synergies (the main branch holds the stand-alone single-cohort tool).

## Acknowledgements

We thank Dr. Krzysztof Mnich and Piotr Stomma from the University of Białystok for their expertise in the Multi-Dimensional Feature Selection algorithm and insightful contributions to our work; Professor Klas Udekwu from the University of Idaho and Dr. Andre Kahles from ETH Zurich for their valuable feedback which substantially improved the quality and clarity of the Manuscript; Dr. Tomasz Kościółek from the Sano Centre for Computational Medicine for inspiring discussions; and the members of Bioinformatics Research Group at MCB JU for their comments and support.

Portions of this text were drafted and structured with the assistance of Claude Opus 4.8 and the methodology was filled in using Claude Fable 5.1. The final version was thoroughly reviewed, verified, and edited by the authors who take full responsibility for the content.

## Supplement Tables

Supplementary Table 1. Seed stability of the MDFS pair selection. For each data type, number of MDFS runs N and consensus threshold k (pair kept if among the per-run top 2% in at least k of N runs): mean number of consensus pairs per LOCO fold and the Jaccard agreement (mean and minimum over the 14 folds) between consensus sets built from two disjoint blocks of seeds (42-51 and 52-61). The pipeline uses N = 10, k = 6.

Supplementary Table 2. The 30 directed synergistic feature pairs with the highest mean joint 2D information gain across the taxa-only, functions-only and combined analyses. base_IG_1D_mean: 1D IG of the base feature; total_IG_2D_mean: joint IG of the pair; ig_difference_raw and ig_difference_pct: IG added by the contributing feature, absolute and relative to the base feature’s 1D IG; pair_ig_count: number of fold-run combinations (of 140 = 14 LOCO folds × 10 MDFS runs) in which the pair was among the top 2% of pairs (pairs selected in fewer than 84 of the 140 fold-runs are included when their mean total IG over those fold-runs ranks among the 30 highest).

Supplementary Table 3. Sensitivity of the LOCO results to the training-set composition: mean and sample-size-weighted AUC of the three feature configurations, with and without the three cohorts that are never held out (AtezoTRIBE, COLOBIOME, NSHII) in the training sets, and the 64 stable pairs shared by the two runs.

Supplementary Table 4. A final set of highly frequent synthetic feature pairs in the CRC data. For each pair, we report the fold selection frequency of the synthetic feature with the frequencies of its two component features. Selection % gain quantifies the improvement of the synthetic pair over its components. Only pairs meeting the selection frequency threshold of >=80% and showing positive gain are included. For each pair the post-selection effect-size summary is given: the transformation evaluated in most cohorts, the number of two-class cohorts, the Benjamini–Hochberg q-value of the Stouffer meta p-value across cohorts, the median rank-biserial correlation and the number of cohorts with positive and negative direction.

Supplementary Table 5. Cohort-adjusted logistic regression for the stable pairs: likelihood-ratio tests of the additive model (cohort + both constituents) against the cohort-only model and of the model with a multiplicative interaction term against the additive model, with Benjamini–Hochberg q-values, and Spearman correlations, on the same pooled samples, between each synthetic feature (the encoding used in Figure 2) and its constituents and between the two constituents.

Supplementary Table 6. Transformation preference of synthetic feature pairs. For every pair selected in >= 50% of LOCO folds as a log-ratio (LR) and/or a geometric mean (GM), the encodings that reached this threshold (Types_Present) and the maximum selection frequency.

Supplementary Table 7. Candidate conserved markers: synergistic pairs identified independently in at least two cohorts with the same diagnosis (only inflammatory bowel disease, HallAB_2017 and NielsenHB_2014, met this criterion). Pairs were matched irrespective of orientation; orientation_per_cohort gives the (base → contributing) direction reported in each cohort and same_orientation_in_all_cohorts whether it agrees between cohorts. IG values are means over the two cohorts; mean_ig_gain_pct is the IG added by the contributing feature relative to the 1D IG of the base feature and is very large when that IG is close to zero (Blautia hydrogenotrophica in HallAB_2017: 0.014).

Supplementary Table 8. Sensitivity of the pair search to the MDFS discretisation (LOCO fold-0 training set, taxonomic features): number of synergistic pairs and overlap with the default setting (Jaccard index of all pairs and of the top-2% pairs; Spearman correlation of the joint IG of shared pairs) when the admissible range of cut points is widened to the one-dimensional default or halved, when three bins per feature are used, and, for reference, when only the random seed is changed.

Supplementary Table 9. Negative controls for the pair search (LOCO fold-0 training sets, seed 42): number of 2D-significant features and of synergistic pairs returned by the pipeline with the real labels and with labels permuted across samples (five permutations).

Supplementary Figure 1. Coverage of feature-selection methods by one another on the LOCO training sets. For each of the 14 LOCO folds and each feature space (taxa, functions, combined), the relevant features were determined with MDFS-2D, MDFS-1D, Welch’s t-test, the Mann–Whitney U test, LASSO logistic regression and ReliefF, and, as a size-matched univariate reference, the N features with the highest 1D IG, where N is the number of MDFS- 2D features of the fold (1D top-N; Methods). Cell (row A, column B) gives the percentage of the features selected by method A that method B also selected, averaged over the folds; n is the mean number of features selected per fold. ReliefF and 1D top-N are given the same number of features as MDFS-2D, so for these two methods the two coverages are identical by construction and measure overlap rather than coverage. MDFS-2D recovers almost all features selected by MDFS-1D and by the two univariate tests, whereas these methods cover only 53–84% of the MDFS-2D features; a feature that is relevant in 1D is also relevant in 2D, so this asymmetry is expected and partly reflects the larger MDFS-2D sets. The size-matched reference shows what the 2D search adds: 81% (taxa), 94% (functions) and 88% (combined) of the MDFS-2D features are also among the N features with the highest 1D IG, and the remaining 46, 24 and 75 features per fold enter the selection because of their pairwise rather than their individual signal. On average 83, 59 and 147 MDFS-2D features per fold (taxa, functions, combined) are selected by none of the three univariate tests, and 23, 4 and 27 by none of the five other selectors. LASSO returns a sparse model whose size varied between folds (combined space: 140–361 features, mean 178); its penalty was chosen by cross-validation with a bounded number of solver iterations (Methods).

Supplementary Figure 2. Per-cohort LOCO-CV performance of univariate feature-only versus synthetic Random Forest models. Scatter plots summarize leave-one-cohort-out cross-validation performance across test cohorts for three feature modes: taxa, functions, and combined. The dashed diagonal represents equal performance between models; points above the diagonal indicate improved performance of the synthetic model relative to the feature-only model for the same cohort, in terms of AUC (a) and accuracy (b).

Supplementary Figure 3. AUC per test cohort (LOCO cross-validation) for taxonomic, functional, and combined models. Grouped bars compare feature-only, synthetic-only, and combined feature sets, highlighting per-cohort variation across feature configurations.

