## Supplementary figures for "Beyond single markers: bacterial synergies identified by Multidimensional Feature Selection reveal conserved microbiome disease signatures"

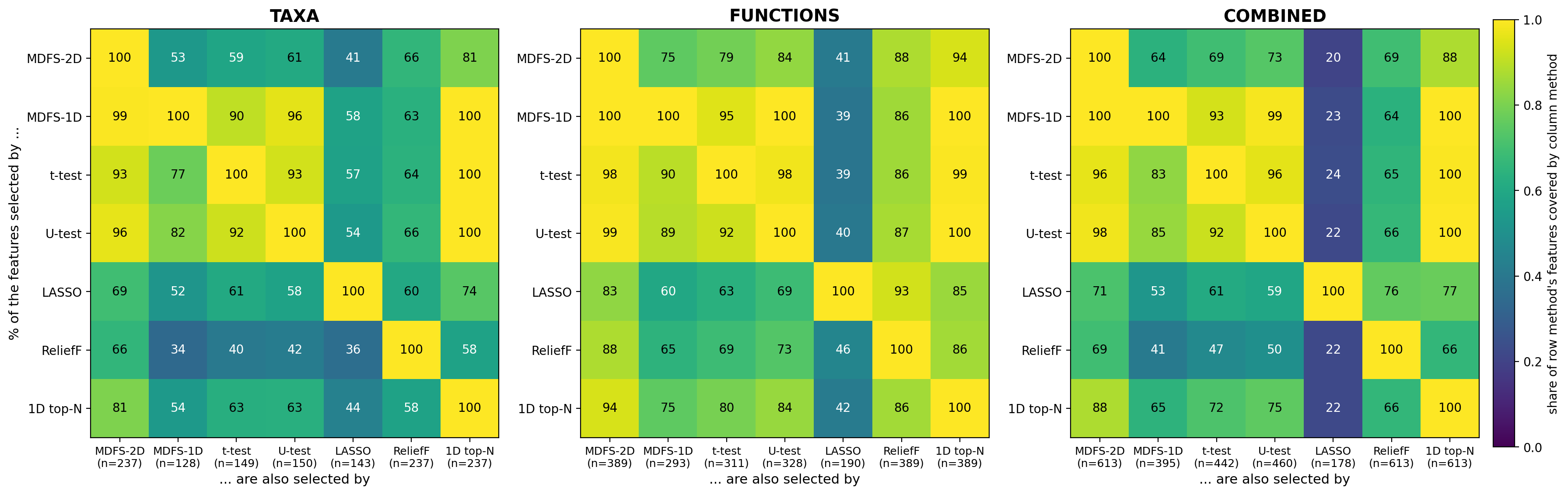

**Supplementary Figure 1.** Coverage of feature-selection methods by one another on the LOCO training sets. For each of the 14 LOCO folds and each feature space (taxa, functions, combined), the relevant features were determined with MDFS-2D, MDFS-1D, Welch's t-test, the Mann–Whitney U test, LASSO logistic regression and ReliefF, and, as a size-matched univariate reference, the N features with the highest 1D IG, where N is the number of MDFS-2D features of the fold (1D top-N; Methods). Cell (row A, column B) gives the percentage of the features selected by method A that method B also selected, averaged over the folds; n is the mean number of features selected per fold. ReliefF and 1D top-N are given the same number of features as MDFS-2D, so for these two methods the two coverages are identical by construction and measure overlap rather than coverage. MDFS-2D recovers almost all features selected by MDFS-1D and by the two univariate tests, whereas these methods cover only 53–84% of the MDFS-2D features; a feature that is relevant in 1D is also relevant in 2D, so this asymmetry is expected and partly reflects the larger MDFS-2D sets. The size-matched reference shows what the 2D search adds: 81% (taxa), 94% (functions) and 88% (combined) of the MDFS-2D features are also among the N features with the highest 1D IG, and the remaining 46, 24 and 75 features per fold enter the selection because of their pairwise rather than their individual signal. On average 83, 59 and 147 MDFS-2D features per fold (taxa, functions, combined) are selected by none of the three univariate tests, and 23, 4 and 27 by none of the five other selectors. LASSO returns a sparse model whose size varied between folds (combined space: 140–361 features, mean 178); its penalty was chosen by cross-validation with a bounded number of solver iterations (Methods).

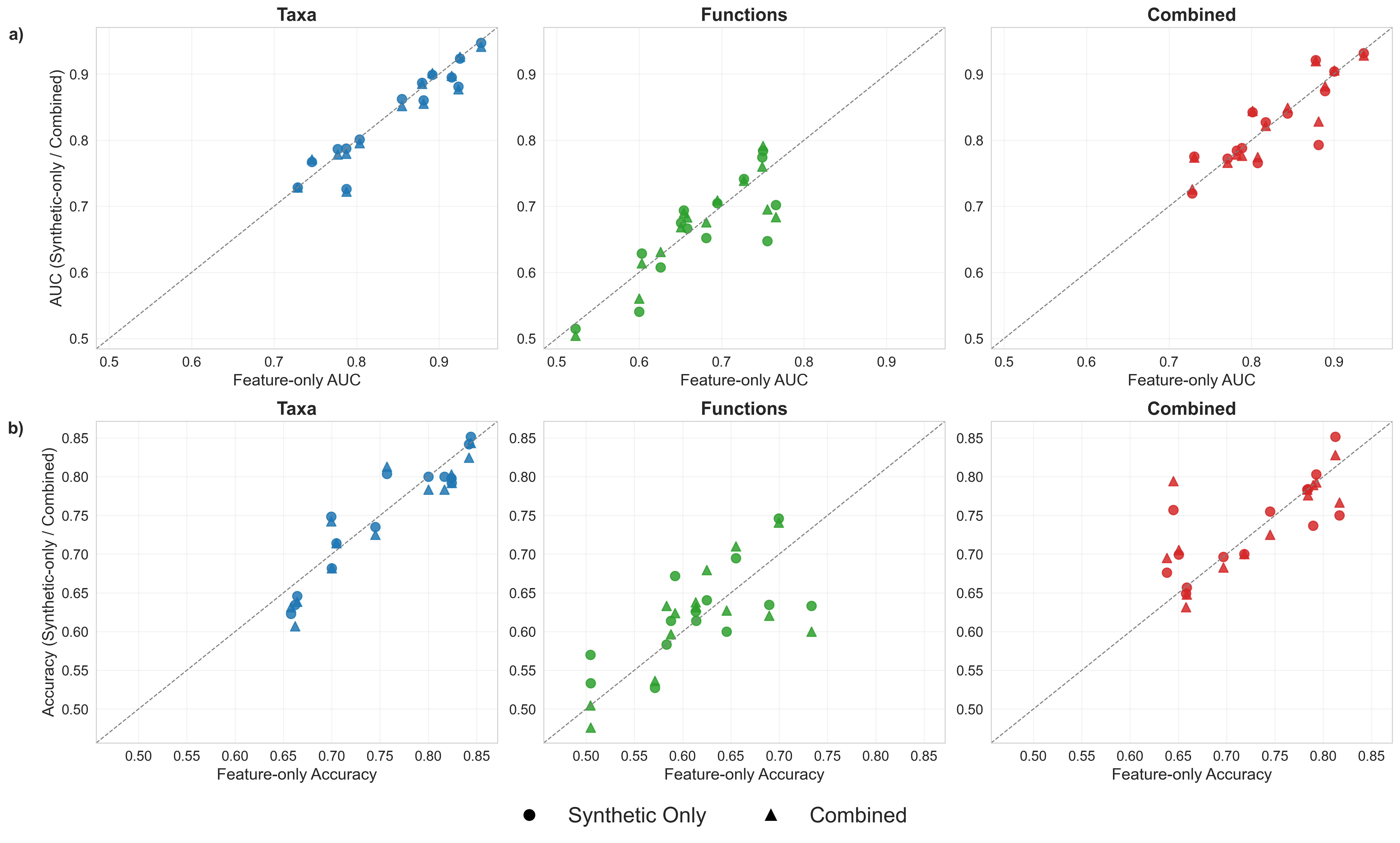

**Supplementary Figure 2.**  Per-cohort LOCO-CV performance of univariate feature-only versus synthetic Random Forest models. Scatter plots summarize leave-one-cohort-out cross-validation performance across test cohorts for three feature modes: taxa, functions, and combined. The dashed diagonal represents equal performance between models; points above the diagonal indicate improved performance of the synthetic model relative to the feature-only model for the same cohort, in terms of AUC (a) and accuracy (b).

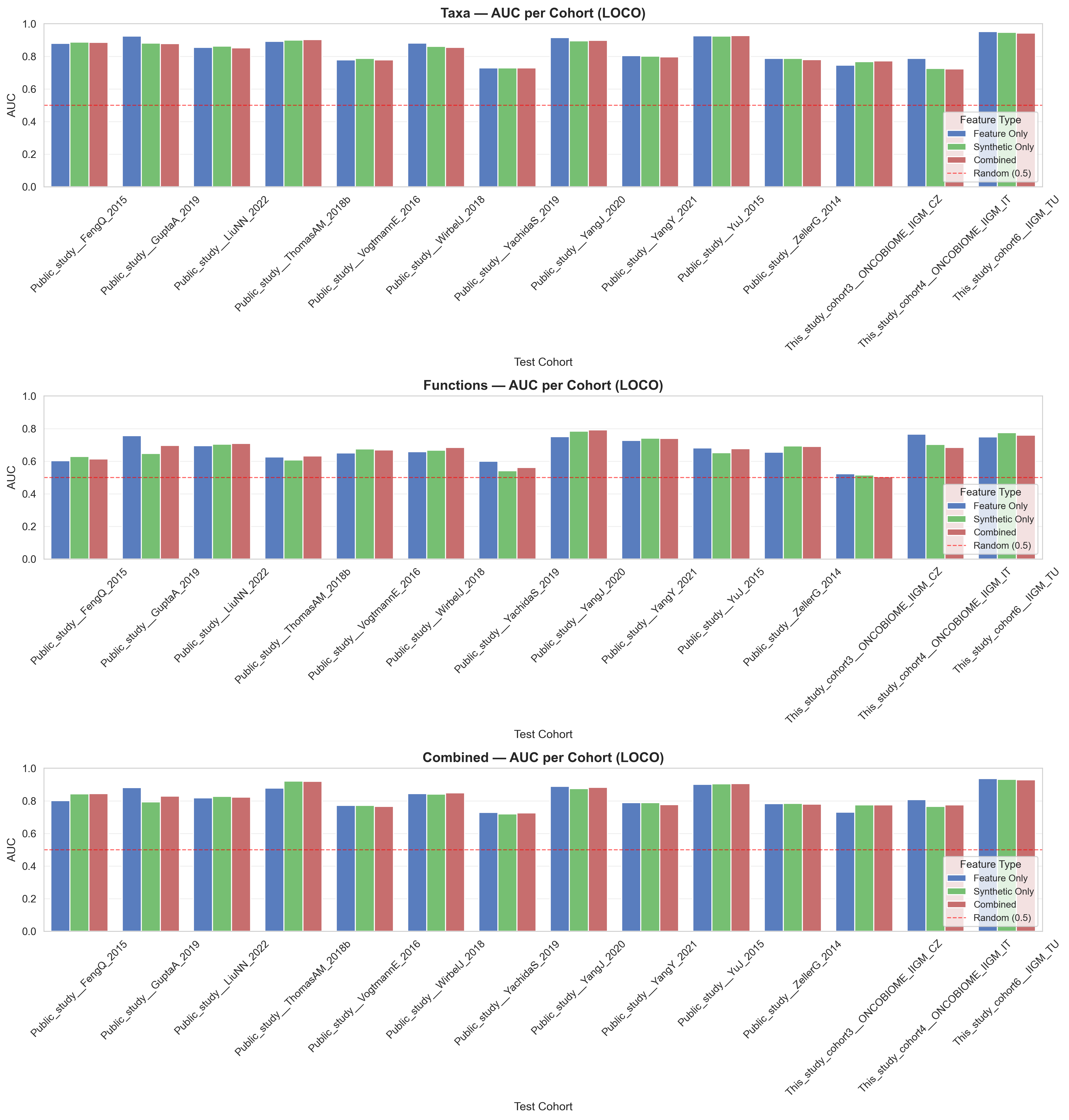

**Supplementary Figure 3.** AUC per test cohort (LOCO cross-validation) for taxonomic, functional, and combined models. Grouped bars compare feature-only, synthetic-only, and combined feature sets, highlighting per-cohort variation across feature configurations.
